# Distinct sensing of BCAAs by mTOR and c-Myc governs T cell proliferation, independent of catabolism

**DOI:** 10.64898/2026.01.16.699967

**Authors:** Kelly Rome, Elise Hall, Aaron Wu, Will Bailis

## Abstract

The branched chain amino acids (BCAAs: leucine, isoleucine, valine) are essential amino acids that function as catabolic substrates and signaling molecules via mTORC1. While individual BCAAs have unique roles in organismal and cellular physiology, the mechanisms underlying their individual effects remain poorly understood. We demonstrate that the three BCAAs have distinct roles in T cell biology. We find that isoleucine and valine are necessary and sufficient for quiescence exit and cell division, whereas leucine is dispensable. Mechanistically, these effects are independent of their diverging catabolic fates and instead due to differential sensing of leucine and isoleucine/valine by mTOR and c-Myc. While isoleucine and valine are necessary and sufficient for c-Myc expression, mTORC1 leucine-sensing represses c-Myc and proliferation during BCAA restriction. Together, we find that the discrete sensing of the BCAAs uncouples two major anabolic regulators, mTORC1 and c-Myc, in cell growth. This provides mechanistic insight into the distinct roles of the BCAAs in cell physiology, highlighting divergent BCAA sensing rather than catabolism, and offering a new lens to appreciate their impact on immunity and pathophysiology.

## INTRODUCTION

The essential BCAAs (Leucine/Leu, Isoleucine/Ile and Valine/Val) have wide-ranging roles in regulating systemic metabolism and metabolic health through their control of cellular anabolism, energy balance and protein translation (Cummings *et al*, 2018; Fontana *et al*, 2016; Neinast *et al*, 2019; Richardson *et al*, 2021; Solon-Biet *et al*, 2019; White *et al*, 2016). In many of these contexts, the BCAAs have been studied as a single metabolic unit due to shared transport and catabolism, however there is a growing body of work demonstrating that the individual BCAAs can exert distinct control over organismal and cellular physiology. Val is uniquely required for proliferation and maintenance of hematopoietic stem cells *in vitro* and *in vivo (Taya et al, 2016)*, as well as for the growth of T cell acute lymphoblastic leukemia (T-ALL) (Thandapani *et al*, 2022). Restriction of dietary Ile and Val, but not Leu, promotes overall metabolic health and improves markers of metabolic disease, and Ile restriction can significantly enhance animal longevity (Green *et al*, 2023; Yu *et al*, 2021). Alternatively, Leu can be sensed and activate the global anabolic regulator mTORC1 through Sestrin2 and the Leu-tRNA synthetase, with Ile acting as a weak mTORC1 agonist (Han *et al*, 2012; Saxton *et al*, 2016; Wolfson *et al*, 2016). In contrast, direct sensors or even direct downstream molecular mechanisms for Val regulated biology remain unknown. These more recent studies support historical observations of individual BCAA antagonism. Excess Leu consumption and subsequent elevation of BCAA catabolism has been shown to restrict animal growth due to systemic Ile and Val depletion, which could be alleviated by Ile/Val supplementation (Calvert *et al*, 1982; Shinnick & Harper, 1977). Collectively, these studies highlight the need to examine the role of individual BCAAs in cell biology.

Amongst primary cells, T cells are well appreciated as a particularly metabolically sensitive lineage. In response to antigen, such as from a tumor or infected cell, T cells must activate, clonally expand and exert immune-promoting functions (ex. cytokine production) to mount an effective immune response. Activated T cells rapidly integrate environmental signals and reprogram nutrient uptake, signaling and metabolism to expand and promote antigen clearance (Chapman *et al*, 2020; Geltink *et al*, 2018; Wilfahrt & Delgoffe, 2024). These rapid metabolic changes are mediated by two master regulators of anabolic growth, mTORC1 and c-Myc (Finlay *et al*, 2012; Shyer *et al*, 2020; Wang *et al*, 2011; Yang & Chi, 2012). Downstream of antigen signaling through the T cell receptor (TCR), these factors promote the expression of System L transporter Slc7a5-mediated intake of large, neutral amino acids, including the BCAAs, which in turn feedback to enforce mTORC1 signaling and c-Myc expression and T cell expansion (Sinclair *et al*, 2013). In this manner, mTORC1, c-Myc, and Slc7a5 form a circuit enforcing anabolic growth. However, the precise molecular mechanisms explaining how each of these components cross-talk and reinforce one another remain a key knowledge gap.

The most well-established mechanism in this pro-growth triad is the requirement of Slc7a5-facilitated Leu uptake for sustained mTORC1 activation. In contrast, the Slc7a5-imported amino acids responsible for c-Myc expression or clonal expansion remain unresolved. Indeed, studies in natural killer (NK) cells have found that Slc7a5-dependent c-Myc induction occurs even in the absence of Leu, suggesting mTORC1 and c-Myc regulation are independently regulated by the transporter (Loftus *et al*, 2018). In addition to their environmental availability, catabolic enzymes regulate intracellular BCAA levels and can modulate mTORC1 signaling in T cells (Ananieva *et al*, 2014). Moreover, systemic alterations in BCAA catabolism have been found to impact T cell immunity *in vivo*, though the molecular mechanisms are unresolved (Yao *et al*, 2023).Together, these studies demonstrate the importance of BCAA utilization in T cell biology. Despite this, the relative contribution of individual BCAAs and BCAA sensing versus catabolism in regulating T cell proliferation or function remains unclear.

Here we identify divergent requirements for each of the BCAAs in governing T cell function, proliferation and pro-growth signaling. We find that Ile and Val are necessary and sufficient for T cell activation, function and cell cycle entry and division, whereas Leu is largely dispensable. This regulation is independent of BCAA catabolism, demonstrating the differential sensing of Leu from that of Ile and Val by nutrient sensitive signaling effectors. To this end we find that Ile and Val are necessary and sufficient for maintenance of c-Myc expression, with their availability leading to rapid changes in protein production. Surprisingly, we demonstrate that this Ile/Val control of c-Myc and cell division is antagonized by Leu sensing by mTORC1, thereby uncoupling mTOR and c-Myc in the regulation of cell growth. These data help clarify the extracellular signals governing proliferation and anabolic growth in T cells, shedding light onto the context-dependent requirements for mTORC1 in T cell biology. Altogether, we offer mechanistic insight into the distinct activities of individual BCAAs beyond their disparate catabolic fates, highlighting how the BCAAs can each govern cell physiology and cell growth through various nutrient-responsive pathways.

## RESULTS

### Leucine is dispensable for T cell cycle entry and proliferation

To investigate the role of each of the BCAAs in T cell biology, we assayed the impact of BCAA restriction on cycle entry, activation, proliferation and function of activated primary CD8 T cells. We observed that BCAA restriction had minimal impact on cell viability during T cell activation, whether T cells were cultured in the absence of one, two or all three BCAAs (**Fig. 1A and B)**. This approach revealed that Val alone was sufficient for upregulation of T cell activation markers CD25 and CD69 (**Fig. 1C**). Interestingly, while BCAAs were required for cell cycle entry, we observed that Leu restriction did not prevent quiescence exit, but rather that Ile and Val alone were sufficient for G1 entry (**Fig. 1D**). To explore whether this activity was specific to initial activation, we activated T cells in complete media before swapping them into varied BCAA-depleted media during days 2-4 post-activation. As with early BCAA restriction and quiescence exit, Ile and Val alone could maintain proliferative capacity and did so in a dose-dependent manner (**Fig. 1E,F**). Extending BCAA-restriction across days 0-3 post-activation (**Fig. EV1A**), we see a similar dose-dependent recovery of cell division only in cells provided with extracellular Ile and Val, whereas by contrast loss of any one BCAA had a detrimental effect on cell viability over the prolonged three day culture period (**Fig. EV1B,C**), consistent with their classification as essential amino acids. Beyond activation, our analysis revealed that extracellular Val is sufficient to support cytokine production, despite the lack of proliferation in these cultures (**Fig. EV1D**). Amino acid restriction has previously been shown to promote macropinocytosis and proteolysis as an alternate mode of amino acid acquisition (Lee *et al*, 2019). To test whether T cells might likewise recover BCAAs through macropinocytosis during restriction to permit cell cycle entry, we treated cells with the macropinocytosis inhibitor EIPA, which inhibits constitutive macropinocytosis in T cells (Charpentier *et al*, 2020). Even in the presence of EIPA, we found that Leu-restricted T cells could still maintain proliferative capacity, with Ile-restricted cells displaying a persistent defect (**Fig. EV1E,F**). Together, these results demonstrate that individual BCAAs each exert discrete effects on activated CD8 T cell biology.

**Figure 1:**
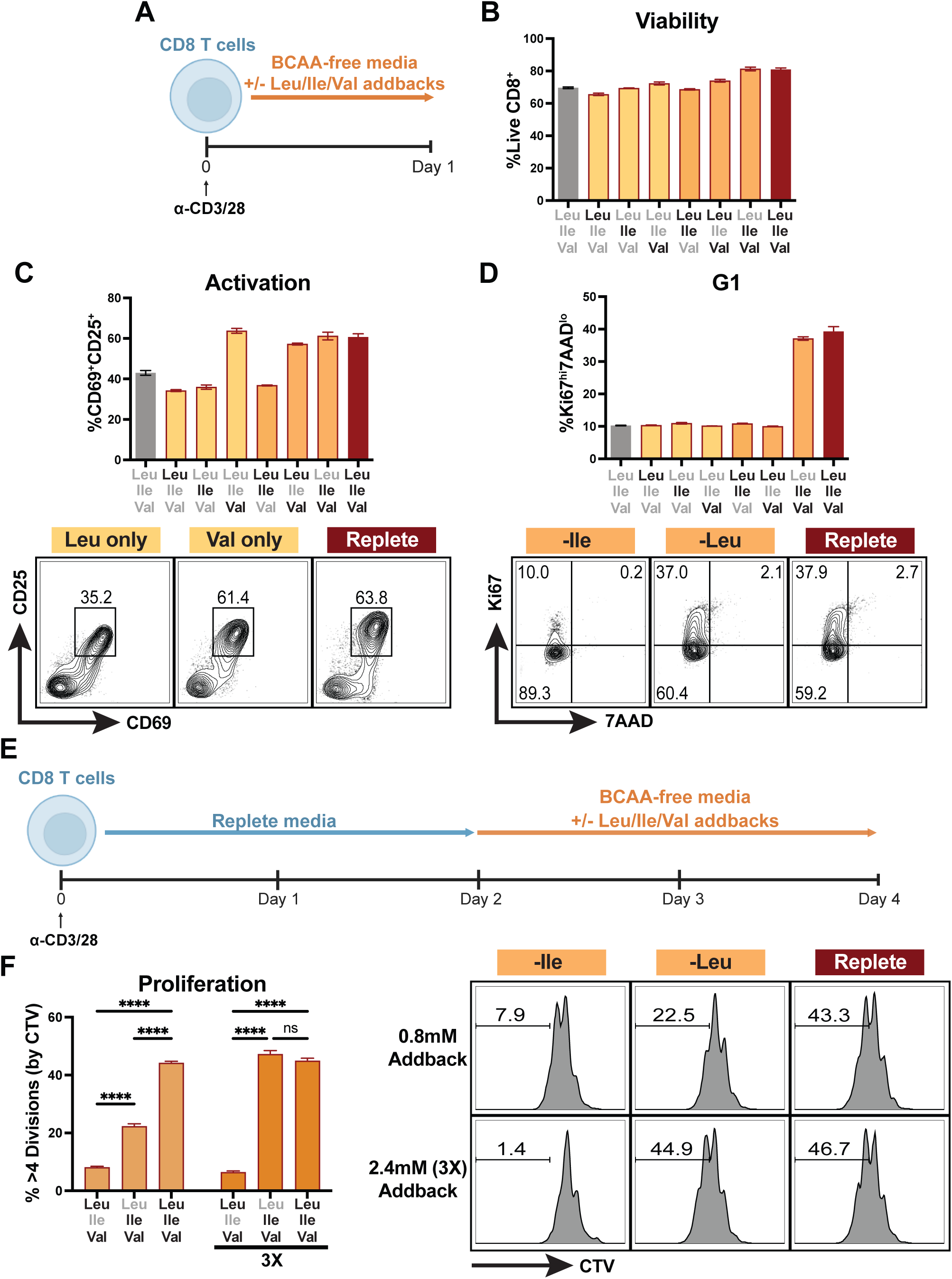
Leucine is dispensable for T cell cycle entry and proliferation. (A) Schematic of Day 0-1 BCAA restriction and addback culture. Bulk CD8 T cells were isolated and purified from spleens of WT mice, activated with anti-CD3 and anti-CD28 antibodies (2ug/mL each) for 1 day in BCAA-free T cell media (TCM) with addback of varying combinations of Leu, Ile and Val (0.8mM each). (B) Frequency of viable CD8 T cells after Day 0-1 BCAA culture. (C,D) Frequency and representative flow plots of T cells expressing activation markers and entering cell cycle after Day 0-1 BCAA culture. (E) Schematic of Day 2-4 BCAA restriction and addback culture. Bulk CD8 T cells were isolated and purified from spleens of WT mice, activated with anti-CD3 and anti-CD28 antibodies (2ug/mL each) for 2 days in replete TCM followed by 2 days in BCAA-free TCM with addback of varying combinations of Leu, Ile and Val (0.8mM or 2.4mM [3X] each as indicated). (F) Frequency and representative flow plots of highly dividing (>4 divisions) cells as measured by Cell Trace Violet (CTV) dilution in Day 2-4 BCAA culture (CTV stain performed immediately after CD8 purification on Day 0 of culture). All error bars are representative of 3 technical replicates. Statistical significance in (F) was calculated using two-way ANOVA with multiple comparisons and Tukey’s correction. ****p<0.0001.

### Isoleucine and valine are necessary and sufficient for c-Myc expression

mTORC1 sensing of Leu – and to a lesser extent Ile – is the most established mechanism for how intracellular BCAAs can differentially regulate cell signaling (Saxton *et al*., 2016; Wolfson *et al*., 2016). Indeed, Leu availability is understood to be required for mTORC1 activation in both CD4 and CD8 T cells (Ananieva *et al*., 2014; Sinclair *et al*., 2013). To assess the direct impact of each BCAA on anabolic signaling, we allowed cells to activate in replete media for one day before swapping into our varied BCAA-restricted culture conditions for 2-4 hours (**Fig. 2A**). Consistent with the literature, we found that Leu, and secondarily Ile availability, promoted mTORC1 signaling in activated CD8 T cells, whereas Val alone could not (**Fig. 2B**). We also assayed induction of the integrated stress response (ISR), a signaling pathway which rewires transcription and translation to recover from environmental stress (Pakos-Zebrucka *et al*, 2016). In the presence of amino acid restriction, the ISR triggers phosphorylation of translation regulator e-IF2α and the selective induction of ATF4 transcription factor translation (Harding *et al*, 2003; Lu *et al*, 2004). We find that acute loss of Leu, Ile or Val leads to differential induction of the ISR (**Fig. EV2**), further supporting the notion that the availability of individual BCAAs is uniquely sensed and integrated by the cell.

**Figure 2:**
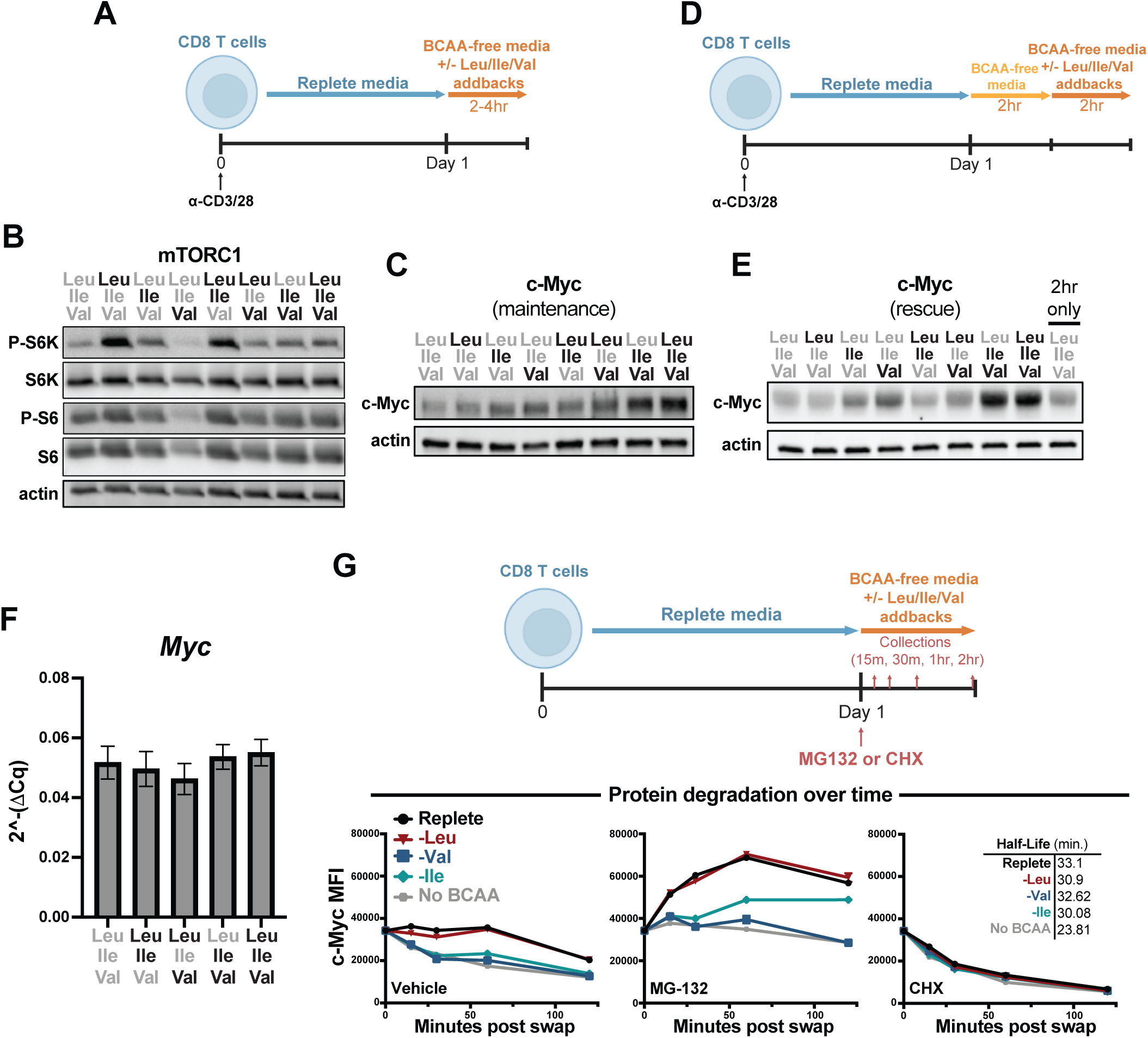
Isoleucine and valine are necessary and sufficient for c-Myc expression. (A) Schematic of acute BCAA restriction and addback culture. Bulk CD8 T cells were isolated and purified from spleens of WT mice, activated with anti-CD3 and anti-CD28 antibodies (2ug/mL each) for 1 day in replete TCM followed by 2-4hr in BCAA-free TCM with addback of varying combinations of Leu, Ile and Val (0.8mM each). (B,C) Western blot for expression/activation of mTORC1 signaling pathway and c-Myc in acute BCAA culture. (D) Schematic of acute BCAA restriction (without addback) and subsequent addback rescue culture. Bulk CD8 T cells were isolated and purified from spleens of WT mice, activated with anti-CD3 and anti-CD28 antibodies (2ug/mL each) for 1 day in replete TCM followed by 2hr in BCAA-free TCM followed by 2hr in BCAA-free TCM with addback of varying combinations of Leu, Ile and Val (0.8mM each). (E) Western blot for c-Myc expression in acute rescue BCAA culture. (F) qPCR for Myc transcript during acute BCAA culture (as in A, 4hr). (G) Schematic and c-Myc expression (by flow cytometry, mean fluorescent intensity) over time following acute BCAA culture (2.4mM addbacks) with addition of cycloheximide (50ug/mL) or MG132 (10uM) at time of BCAA culture and with collections at 15, 30, 60 and 120 minutes following drug treatment/BCAA restriction and addback. All error bars are representative of 3 technical replicates. Half-lives in (G) calculated using one phase decay non-linear regression analysis.

The large neutral amino acid (LNAA) transporter Slc7a5, which transports the BCAAs among other amino acids into a cell, is required for T cell expansion and c-Myc expression following antigen-stimulation (Sinclair *et al*., 2013). Given this and our data demonstrating a critical role for intracellular BCAAs in regulating cell cycle entry and proliferation, we investigated how acute restriction of the individual BCAAs affects c-Myc expression in activated CD8 T cells. Consistent with their role in proliferation, we found that Ile and Val are sufficient to maintain c-Myc expression following T cell activation (**Fig. 2C**). To determine if the effect of BCAA restriction on c-Myc expression was reversible and rapidly sensitive to Val and Ile availability, we cultured cells at day one post-activation for two hours in BCAA-depleted culture media and then reintroduced BCAAs for an additional two hours (**Fig. 2D**). We found that provision of Ile and Val could directly induce expression of c-Myc comparably to BCAA replete culture media, and evidence that Leu could be antagonistic to c-Myc expression during nutrient stress (**Fig. 2E**). We then sought to determine the mode of regulation through which the BCAAs regulated c-Myc expression. To do so, we first examined the impact of BCAA restriction on *Myc* transcript levels and observed minimal impact in any condition (**Fig. 2F)**. As c-Myc protein synthesis and stability is understood to be the primary mechanism for regulation, we next assessed how BCAA availability affected these readouts. We find that Leu-restriction did not alter c-Myc protein stability nor the rate of protein accumulation in the absence of proteasome activity (**Fig. 2G**). In contrast, Ile and Val restriction each impaired the rate of c-Myc accumulation in the presence of MG132, while minimally impacting c-Myc half-life, suggesting a role for Val and Ile in supporting c-Myc translation. Collectively, these data identify an essential role for Ile and Val sensing in the induction of pro-proliferative gene expression.

### BCAA-specific effects on T cell biology are independent of their catabolism

Upon entering a cell, the BCAAs can enter the mitochondria as their respective α-ketoacids, where they are irreversibly decarboxylated and committed to catabolism by the branched chain α-ketoacid dehydrogenase complex (BCKDC) (**Fig. 3A**) (Ichihara & Koyama, 1966; Johnson & Connelly, 1972; Neinast *et al*., 2019). Downstream of the BCKDC, metabolism diverges between the individual BCAAs, particularly for Leu compared to Ile and Val, and thus individual BCAA-specific metabolism could account for the distinct requirements we observe for Ile and Val in T cell biology. To test this, we crossed CD4-cre mice to *Dbt*-floxed animals to specifically ablate *Dbt* expression in T cells (“Dbt cKO”, **Fig. 3A**). Dbt is one of four components of the BCKDC, all of which are required for the complex to function (Neinast *et al*., 2019), and thus Dbt cKO T cells have lost the capacity to metabolize BCAAs. Using this system, we identified that BCAA catabolism is dispensable for cell cycle entry, c-Myc expression or proliferation in both BCAA-restricted or replete culture media (**Fig. 3B-E**). We also found no changes in activation markers or cell viability at one day post-activation (**Fig. EV3A,B**). These data suggest that while BCAAs, notably Ile and Val, are required for T cell activation, c-Myc expression, cell cycle entry and proliferation, these effects occur independently of BCAA catabolism and instead are explained by alternate mechanisms.

**Figure 3:**
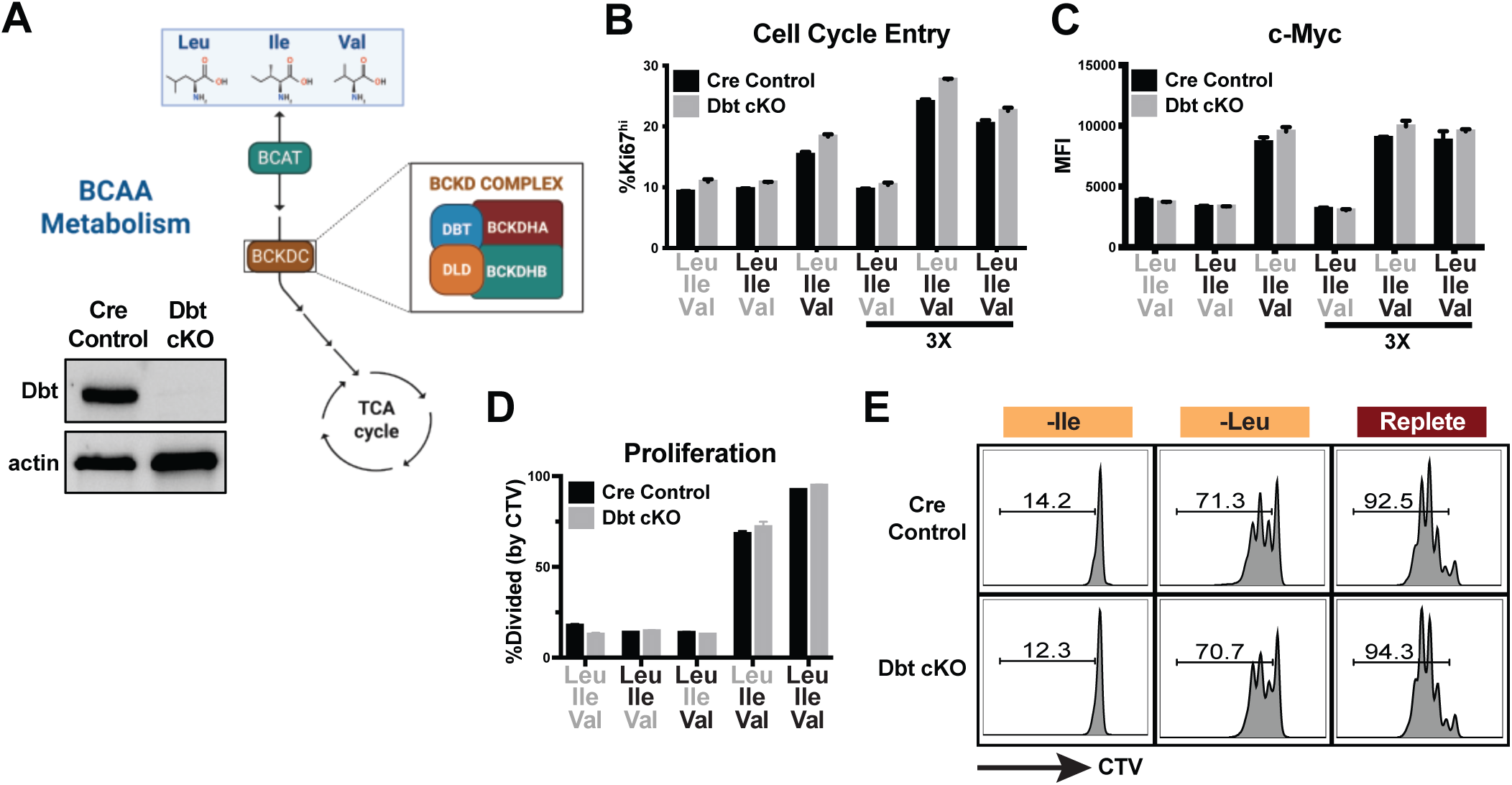
BCAA-specific effects on T cell biology are independent of their catabolism. (A) Schematic of BCAA catabolism and ablation of Dbt expression (Western blot) in purified CD8 T cells from Dbt conditional knockout (CD4-cre) mice. (B,C) Frequency of cells entering cell cycle and expression of c-Myc (flow cytometry, MFI) after Day 0-1 BCAA culture as described in (1A) (0.8mM or 2.4mM [3X] BCAA addbacks as indicated). (D,E) Frequency and representative flow plots of dividing cells (% of cells with CTV dilution, CTV stain performed on Day 0) in Day 0-3 BCAA culture (2.4mM addbacks) as described in (EV1A). All error bars are representative of 3 technical replicates.

### BCAA restriction uncouples mTORC1 and c-Myc in CD8 T cell proliferation

Having found distinct, catabolism-independent roles for Leu and Ile/Val in regulating CD8 T cell proliferation, we next hypothesized that differences in BCAA sensing might be responsible for the disparate phenotypes observed. Indeed, we had found that Leu is dispensable for and potentially antagonistic to c-Myc expression during nutrient restriction (Fig. 2E). We therefore tested whether sensing of Leu by mTOR might contribute to this antagonism, as well as the potential role of the ISR in detecting amino acid stress. To do so, we subjected CD8 T cells to acute BCAA restriction in the presence or absence of the mTOR inhibitor rapamycin or ISRIB (**Fig 4A**), a drug that relieves the effects of eIF2α phosphorylation on translation. While ISRIB treatment suppressed ATF4 expression as expected, it had minimal impact on c-Myc protein levels (**Fig 4B**). In contrast, mTOR inhibition resulted in the full or partial restoration of c-Myc expression in Ile and Val deprived CD8 T cells (**Fig. 4B**). We next tested whether recovery of c-Myc expression corresponded to a similar recovery in proliferative capacity. We found that mTORC1 inhibition was sufficient to enhance T cell proliferation during Ile withdrawal, whereas it had minimal to slightly detrimental impacts on proliferation in Leu-restricted or replete culture media (**Fig. 4C-E**). We further observed that suppressing mTORC1 activity promoted S-phase progression in Ile-restricted cells, as determined by Edu incorporation, while failing to affect Leu-restricted or control cells, suggesting that Ile/Val are required for optimal cell cycle progression kinetics (**Fig 4F**). To explore the potential mechanism underlying this, we examined how BCAA-restriction and mTORC1 activity differentially affected the negative cell cycle regulators p21 and Rb. We observed that mTORC1 inhibition resulted in the loss of p21 accumulation and the restoration of Rb phosphorylation in BCAA-restricted cells, indicating a relief in cell cycle arrest (**Fig. 4G**). Given that mTORC1 is understood to be required for T cell quiescence exit rather than sustained proliferation, we next aimed to test if this antagonistic role for mTORC1 on the cell cycle during BCAA-restriction was unique to actively cycling T cells or if it could also be revealed at the time of activation. To test this, we repeated the restriction and mTORC1 inhibition assay over day 0-1 of T cell activation (**Fig. EV4A**). In contrast to our observations in later-stage activated T cells, we observed that mTORC1 signaling was still required for quiescence exit in both control and Leu-restricted conditions, while failing to restore cell cycle entry in Ile or Val deprived cells (**Fig. EV4B**). Thus, mTORC1 plays distinct roles in regulating the cell cycle during quiescence exit versus once cells have already entered. At early time points, it behaves as a canonical pro-growth effector alongside c-Myc to promote cell cycle entry. Once T cells have entered the cell cycle however, we find that BCAA-restriction uncouples mTORC1 and c-Myc in regulating CD8 T cell proliferation, with mTORC1 antagonizing the cell cycle. Altogether, our data support a model in which the BCAAs act through distinct axes of anabolic regulation, a Leu-mTORC1 axis and an Ile/Val-c-Myc axis, which become uncoupled upon BCAA restriction, shedding mechanistic light on the distinct effects individual BCAAs exert on cell biology (**Fig. 4H**).

**Figure 4:**
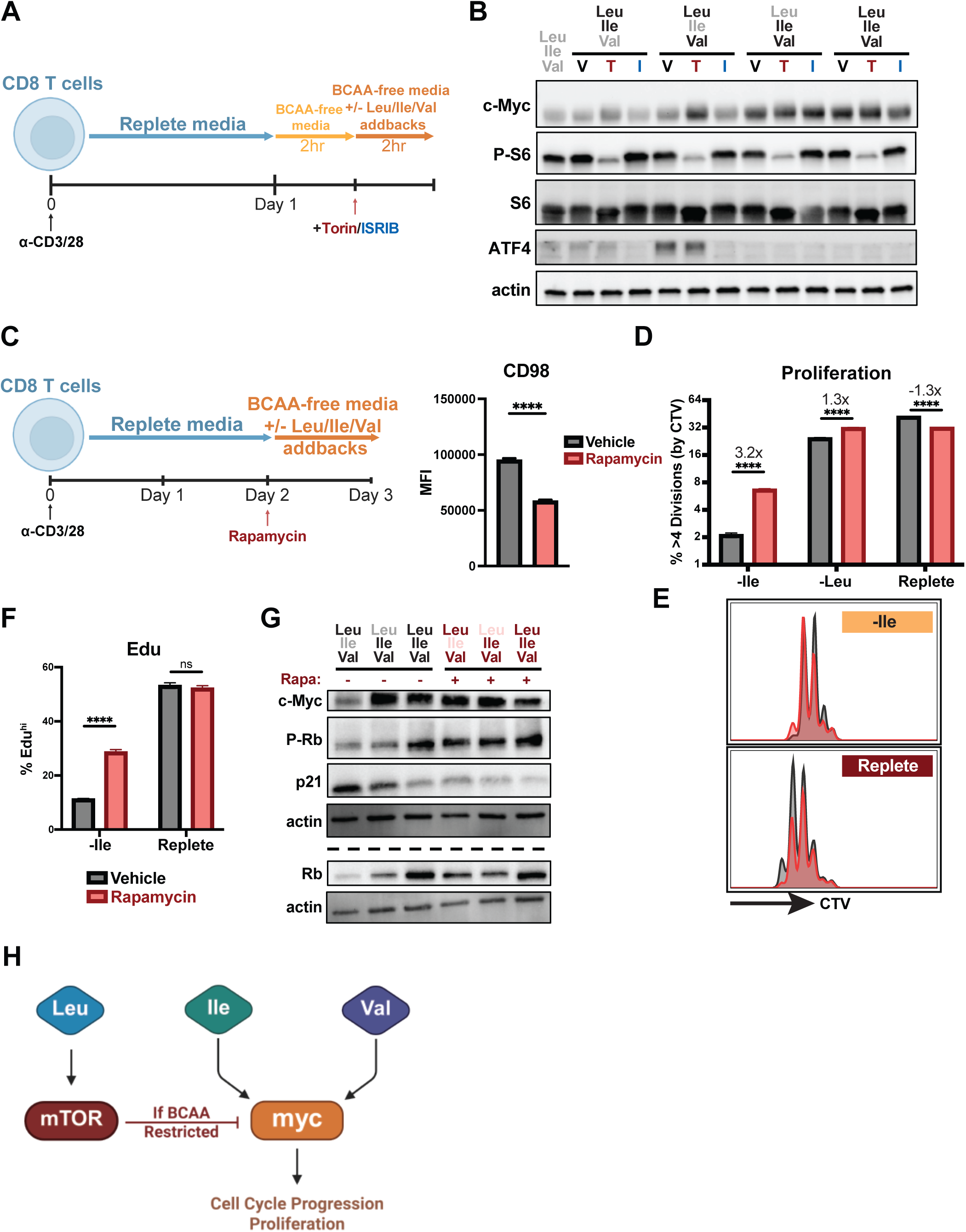
BCAA restriction uncouples mTORC1 and c-Myc in CD8 T cell proliferation. (A) Schematic of acute BCAA restriction (without addback) and subsequent addback rescue culture with Torin/ISRIB treatment. Bulk CD8 T cells were isolated and purified from spleens of WT mice, activated with anti-CD3 and anti-CD28 antibodies (2ug/mL each) for 1 day in replete TCM followed by 2hr in BCAA-free TCM followed by 2hr in BCAA-free TCM with addback of varying combinations of Leu, Ile and Val (0.8mM each) as well as treatment with ISRIB (1uM), Torin (1uM) or vehicle control. (B) Western blot for expression of c-Myc as well as confirmation of ISR inhibition (ATF4 expression) and mTOR inhibition (phosphorylation of S6) in acute BCAA rescue culture with Torin/ISRIB. (C) Schematic of Day 2-3 BCAA restriction with rapamycin and addback culture. Bulk CD8 T cells were isolated and purified from spleens of WT mice, activated with anti-CD3 and anti-CD28 antibodies (2ug/mL each) for 2 days in replete TCM followed by 1 day in BCAA-free TCM with addback of varying combinations of Leu, Ile and Val (0.8mM each) and simultaneous treatment with 50nM rapamycin or vehicle control. Validation of mTOR inhibition by rapamycin in complete media control cultures via expression of CD98 (mTOR target). (D,E) Frequency and representative flow plot of highly dividing (>4 divisions) cells as measured by CTV dilution in Day 2-3 BCAA culture with rapamycin (CTV stain performed immediately after CD8 purification on Day 0 of culture). (F) Frequency of cells incorporating Edu in Day 2-3 BCAA culture with rapamycin (addition of 10uM Edu during last 4 hours of culture). (G) Western blot for expression/activation of cell cycle regulators (p21, p/Rb) and c-Myc after Day 2-3 BCAA culture with rapamycin. (H) Schematic of proposed model representing how Leu, Ile and Val sensing uncouples mTORC1 and c-Myc regulation to govern T cell proliferation. All error bars are representative of 3 technical replicates. Statistical significance in (C) was calculated using Welch’s t test. Statistical significance in (D,F) was calculated using two-way ANOVA with multiple comparisons and Sidak’s correction. ****p<0.0001.

## DISCUSSION

The BCAAs are essential nutrients that can both fuel catabolism and tune anabolic signaling, underlying physiological processes ranging from muscle function to metabolic health and longevity. Despite sharing transport and catabolic pathways, there is growing evidence that the individual BCAAs play distinct roles in organismal and cellular biology. Ile and Val for example are distinctly required for regulating stem cell maintenance as well as metabolic disease and life-span (Green *et al*., 2023; Taya *et al*., 2016; Yu *et al*., 2021). However, the molecular mechanisms underlying the disparate activities of Leu, Ile and Val have proven elusive.

Here we explore the distinct impact each BCAA has on the cellular biology of T cells. We demonstrate that T cells have distinct requirements for Ile and Val in cell growth and function, independent from BCAA catabolism. Mechanistically, we show that Ile and Val are necessary and sufficient for maintaining c-Myc expression in activated T cells, while Leu and mTORC1 signaling antagonize both c-Myc expression and cell division during nutrient stress. Altogether we reveal distinct, catabolism independent roles for individual BCAAs through their differential regulation of two key anabolic regulators, c-Myc and mTORC1, shedding new light on how Ile and Val regulate cell growth and proliferative signaling.

c-Myc and mTOR are central pathways governing anabolic growth. Both mTOR and c-Myc induce expression of amino acid transporters such as Slc7a5, promoting amino acid import that reinforces their own activation creating a feedforward loop enforcing cell expansion (Nachef *et al*, 2021; Park *et al*, 2017; Yue *et al*, 2017). While this relationship between mTOR, c-Myc, and general amino acid transport is well substantiated, how the import of specific amino acids informs this network is incompletely understood. mTORC1 activation and sensing by Leu is one of the clearest examples of individual BCAAs exerting distinct impacts on cell signaling, as we observe in our study. However, key studies in T and NK cells have found that sustained c-Myc expression downstream of Slc7a5 occurs independently of Leu availability and mTORC1 activity, suggesting parallel regulation of these two pathways (Loftus *et al*., 2018; Sinclair *et al*., 2013). Here we show that beyond Leu-mTORC1 interactions, Ile and Val directly promote c-Myc expression and cell division following T cell activation in the absence of extracellular Leu. Not only do we observe a dichotomy in how mTORC1 and c-Myc respond to each of the BCAAs, we further find Leu and mTORC1 activation antagonizes the ability of Ile and Val to stabilize c-Myc expression during amino acid restriction. These findings help address a longstanding knowledge gap in how mTORC1 and c-Myc activity are orchestrated downstream of amino acid transport.

Dividing cells require rapid integration of extracellular signals and nutrient sensing to orchestrate cellular growth. In the context of cancer, mTOR and c-Myc signaling often enforce the activity of each other, or integrate in parallel to facilitate cancer cell growth and survival, as is further evidenced by the synthetic lethality of mTOR inhibition in many settings of c-Myc driven oncogenesis (Grzes *et al*, 2017; Liu *et al*, 2017; Mahauad-Fernandez *et al*, 2025; Potts *et al*, 2025; Pourdehnad *et al*, 2013). In activated CD8 T cells specifically, mTOR activation and c-Myc expression is highly correlated, though direct feedback or regulation between the two pathways during expansion or in varied nutrient settings has not been explored (Pollizzi *et al*, 2016). Surprisingly, we find that individual BCAA sensing uncouples these core anabolic signaling processes, as well as demonstrates direct antagonism of these pathways in certain contexts of nutrient stress. While distinct in mechanism, a similar phenomenon has been reported in the context of myeloid development, where, mTOR inhibition leads to elevated c-Myc transcription, suggesting this may be a broader axis of regulation (Lee *et al*, 2017). Our data indicate the most likely mechanism for this regulation in T cells is at the level of c-Myc mRNA translation, with protein stability and transcription unaffected, but rate of production sensitive to Ile and Val availability. Notably, we find that even though BCAA restricted cells engage the ISR, that relieving the effects of eIF2α phosphorylation on translation with ISRIB was not sufficient to restore c-Myc expression. This distinguishes c-Myc from canonical stress-responsive transcripts and suggests that Ile/Val-sensing can function through alternate mechanisms. It is possible that mTOR inhibition restores proliferative capacity and c-Myc protein by globally dampening metabolic and translational activity during amino acid restriction, relieving cellular and ribosomal stress. This parallels observations in lymphoma, where ISR-mediated mTOR suppression promotes c-Myc-driven cell survival by relieving proliferative stress (Tameire *et al*, 2019). However, our data in early quiescence exit or the partial rescue of growth by rapamycin, also indicate that the exacerbation of nutrient stress by Leu-mTORC1-driven metabolic and translational demand may not be the only mechanism at play. Studies have also shown that Leu restriction and mTOR inhibition can both induce macropinocytosis to support amino acid recovery during nutrient restriction and that this can restore cancer cell growth (King *et al*, 2020; Palm *et al*, 2015). In contrast, we show that growth during Leu restriction occurs even when we inhibit macropinocytosis, suggesting that Ile/Val sufficiency in cell division is unlikely due to elevated amino acid recovery induced by Leu-restriction. Instead, these data further support a model of direct sensing of Ile and Val to facilitate pro-proliferative gene expression and division. Future work will be needed to deconvolute both layers of cross-talk underlying mTOR antagonism of c-Myc and identify the Ile and Val specific sensors responsible for their distinct role in this network.

Intriguingly, we find that mTOR-Myc antagonism is context dependent during the course of activation. During the earliest stages of T cell activation, cells undergo major metabolic reprogramming, inducing nutrient transporters, glycolytic enzymes, and ribosome biogenesis. Both mTOR and c-Myc are known to cooperatively build the cellular machinery necessary for expansion (Marchingo *et al*, 2020; Sinclair *et al*., 2013; Tan *et al*, 2017). In keeping with this, we find that mTOR signaling is essential regardless of BCAA restriction as T cells are exiting quiescence. Even in the absence of Leu, T cells remain sensitive to mTOR inhibition, underlining the importance of this pathway at this stage. However, once metabolic reprogramming is established and cells are proliferating (days 2-4), the cellular context changes and we find the relationship between mTOR and c-Myc uncouples during nutrient stress. At this stage, we find the Ile/Val-c-Myc axis can operate independently of mTOR, while active mTOR may impose translational demands that exacerbate nutrient limitation. These findings are in keeping with what is known about the distinct metabolic demands T cell display at different stages of activation and we have observed in T cells broadly responding to nutrient stress in other studies (Bailis *et al*, 2019; Chang *et al*, 2013; Chapman *et al*., 2020; Ma *et al*, 2024; Turner *et al*, 2024; Wilfahrt & Delgoffe, 2024). We have recently shown that CD8 T cells possess a high level of biosynthetic plasticity that allows them to remain resilient to environmental stresses like those found within tumors (Scaglione *et al*, 2025). Altogether, our work supports a model in which CD8 T cells can sustain their core programming through discrete modules that vary in their nutrient sensitivity. Given a T cell is ignorant of the pathogenic or tissue context its antigen will be present, this confers them with the capacity to maintain host fitness across a wide range of potential environmental settings. Understanding how T cells coordinate environmental signals to regulate these pathways has the potential to open new opportunities for therapeutically tuning T cell activity during cancer, chronic infection, autoimmunity, and transplant. For example, rapamycin-sensitivity in various tumor settings may provide a unique opportunity to prioritize exploration of mTOR inhibition in T cell-based therapies, which our data suggest could enhance tumor microenvironment resiliency in T cells and synergize with the rapamycin-mediated suppression of cancer cell growth in sensitive tumors.

In summary, we have identified critical requirements for Ile and Val sensing in cell growth, independent of their catabolism, providing molecular insight into the distinct biology of the BCAAs. We further reveal the selective uncoupling of c-Myc and mTOR by each of the BCAAs during amino acid stress. Our work highlights how anabolic signaling modules, otherwise understood to cooperatively promote cell growth, are differentially regulated by discrete nutrient inputs, enabling cells to adapt their biology to their environment and sustain function. Combined, these data offer a new framework to appreciate the unique roles individual BCAAs play in cellular and organismal physiology. This context-dependent antagonism of Leu-mTOR and Ile/Val-c-Myc axes offers new strategies for targeting T cells in disease and for understanding at the cellular and molecular level how dietary BCAAs impact metabolic health and immunity.

## METHODS

### Mice

C57BL/6J (Jax #000664), CD4-cre (Jax #017336), and OT-I (Jax #003831) mice were obtained from The Jackson Laboratory. Dbt-floxed mice were a gift from Rebecca Ahrens-Nicklas (Kuhs *et al*, 2025). All mice were maintained under specific pathogen-free conditions at the Children’s Hospital of Philadelphia (CHOP). Experiments were performed with male or female mice aged between 6 and 10 weeks of age. OT-I mice were used to isolate larger CD8 T cell numbers as needed. All experiments were performed in accordance with the Institutional Animal Care and Use Committee of CHOP (IAC 24-001325).

### CD8 T cell Isolation and Culture

Murine CD8 T cells were isolated from spleens, following ACK lysis, by negative selection using the EasySep Mouse Isolation kit (StemCell Technologies, 19853). C57BL/6 or OT-I CD8 T cells were stimulated with plate-bound anti-CD3ε (2ug/mL, clone 145-2C11, BioLegend), anti-CD28 (2ug/mL, clone 37.51), and rIL-2 (5ng/mL, BioLegend).

T cells were cultured in Roswell Park Memorial Institute (RPMI) 1640 Medium (Gibco, #11875093), supplemented with 10% dialyzed fetal bovine serum, additional 2mM L-glutamine (Gibco, #25030081), 1mM sodium pyruvate (Gibco, #11360070), 25mM HEPES (Gibco, #15630080), 1x penicillin/streptomycin (Gibco, #15140122), 55uM 2-mercaptoethanol (Gibco, #21985023). For BCAA deprivation media experiments, T cells were cultured in BCAA-free RPMI 1640 Medium w/ L-Glutamine, w/o L-Isoleucine, L-Leucine, L-Valine (US Biological, #R8999-20) and supplemented as above. BCAAs were added back as indicated with Leucine (Alfa Aesar, J62824), Isoleucine (Alfa Aesar, J63045), or Valine (Alfa Aesar, J62943) powder. All cells were cultured in a CO2 incubator at 37C at 5% CO2.

For all experiments using pharmacological agents, the following doses were used unless otherwise noted: EIPA (25uM, MedChem Express), Cycloheximide (50ug/mL, Sigma), MG132 (10uM, Sigma), Torin (1uM, Cayman), ISRIB (1uM, Ambeed), Rapamycin (50nM, Sigma).

### Flow Cytometry

All flow cytometry was performed on a CytoFLEX LX or CytoFLEX S cytometer (Beckman Coulter). Intracellular cytokine production was measured by adding brefeldin A (Invivogen) 30 min after stimulation with 50ng/mL PMA (Thermo) and 500ng/mL ionomycin (Cayman) and culturing for 4-5 hours before surface staining. Cell division was measured by labeling cells with CellTrace Violet (Invitrogen) per manufacturer’s protocols before activation and evaluated for proliferation at the indicated time points. Staining was performed with combinations of the following antibodies: anti-CD8 (1:300, 53-6.7, Thermo), anti-CD69 (1:300, H1.2F3, Thermo), anti-CD25 (1:300, PC61.5, Thermo), Ki-67 (1:600, SolA15, Thermo), anti-INFy (1:200, XMG1.2, Thermo), anti-TNFa (1:200, MP6-XT22, Thermo), anti-c-Myc (1:400, D84C12, Cell Signaling), anti-CD98 (1:300, RL388, Thermo). DNA content was measured by DAPI or 7AAD for cell cycle analyses. Viability was assessed using eBioscience Fixable Viability Dye eFluor 780 (Thermo). For intracellular staining, cells were fixed, permeabilized, and stained for 30min-1hr using the eBioscience™ Foxp3 / Transcription Factor Staining Buffer Set (Invitrogen, #00-5523-00). Samples were run in technical triplicates.

For Edu incorporation assay, cells were cultured as indicated and labeled with 10mM Edu during the final 4 hours of culture. Edu incorporation was detected using the Click-iT Edu Cell Proliferation Kit (AF488, Invitrogen, C10499) according to the manufacturer’s instructions and read out by flow cytometry.

### RT-qPCR

RNA was isolated with the Zymo Quick-RNA MicroPrep Kit (Zymo Research, #R1050) with DNase I treatment according to the manufacturer’s instructions. cDNA was synthesized from purified RNA using Thermo Maxima H Minus Reverse Transcriptase (Thermo, #EP0751) according to the manufacturer’s instructions with 25 pmol each oligo(dT)18 and random hexamer primers using the following thermocycler protocol: 25°C for 10min, 50°C for 15min., 85°C for 5min. cDNA was quantified by real-time quantitative polymerase chain reaction using PowerTrack^TM^ SYBR Green Master Mix (Thermo, #A46110) on a Bio-Rad CFX384 instrument. Technical triplicates were quantified using 2-ΔCt normalized to Rps18. Primers: Mouse Myc F (GTACCTCGTCCGATTCCACG), Mouse Myc R (GCTCTTCTTCAGAGTCGCTGC).

### Western Blot

Cells were pelleted and lysed in ice cold RIPA lysis buffer (50mM Tris pH8.0, 1mM EDTA, 150mM NaCl, 1% NP-40, 0.5% Na-deoxycholate, 0.1% SDS) supplemented with 10mM NaF, 1mM Na3VO4, and cOmplete^TM^ Mini EDTA-free Protease Inhibitor Cocktail (Roche, #11838170001). Protein concentration was quantified by Bradford Assay (Bio-Rad, #5000006) using bovine serum albumin as a standard. 10-20ug of protein was combined with reducing sample buffer (Boston BioProducts, #BP-111R) and run on SDS-PAGE AnykD^TM^ Mini-PROTEAN TGX^TM^ Precast Protein Gels (BioRad, #456-8126) followed by transfer using the Trans-Blot Turbo Transfer System (BioRad). Membranes were blocked with 5% nonfat milk in tris-buffered saline with 0.1% Tween 20 (TBS-T) for 30 minutes-1 hour. Primary antibodies were diluted in TBS + 1% Casein (BioRad, #1610782) and incubated with membranes overnight at 4°C with rocking. Four washes with TBS-T (5-10 minutes each with rocking) were performed before and after a 1 hour secondary antibody incubation (diluted in 5% nonfat milk in TBS-T) with rocking. Membranes were visualized using SuperSignal West Pico PLUS or Femto Chemiluminescent Substrate (Thermo, #34580 and 34095) on a ChemiDoc MP Imager (BioRad). Actin was quantified after incubation with anti-actin fluorescently conjugated antibody for 1 hour at room temperature with rocking. Antibodies used: Phospho-S6K (1:2000, 108D2 Thr^389^, Cell Signaling), S6K (1:1000, 49D7, Cell Signaling), Phospho-S6 (1:5000, D57.2.2E Ser^235/236^, Cell Signaling), S6 (1:2000, 5G10, Cell Signaling), c-Myc (1:1000, D84C12, Cell Signaling), Phospho-eIF2α (1:1000, D9G8 Ser^51^, Cell Signaling), eIF2α (1:1000, L57A5, Cell Signaling), ATF4 (1:1000, D4B8, Cell Signaling), Dbt (1:1000, ProteinTech #12451-1-AP), Phospho-Rb (1:1000, D20B12 Ser^807/811^, Cell Signaling), Rb (1:1000, D20, Cell Signaling), p21 (1:1000, XX118, BD), Actin-hFAB rhodamine (1:2000, BioRad).

### Statistical Analysis

Statistical significance of our findings were calculated with the tests indicated in each Figure Legend using GraphPad Prism software. P values of <0.05 were considered to be statistically significant. All error bars represent the standard error of the mean (SEM). All experiments were repeated across two or more independent experiments.

## ACKNOWLEDGMENTS

We thank the Children’s Hospital Flow Cytometry Core for providing support and instrumentation; Zoltan Arany for feedback and thoughtful discussion; and all members of the Bailis Laboratory for feedback and support. Figure schematics were created in https://BioRender.com.

## AUTHOR CONTRIBUTIONS

Conceptualization: KR, WB

Methodology: KR, EH, WB

Investigation: KR, EH, AW

Visualization: KR

Funding Acquisition: WB

Project Administration: WB

Supervision: WB

Writing – original draft: KR, WB

Writing – review & editing: KR, EH, WB

---

**Figure EV1:**
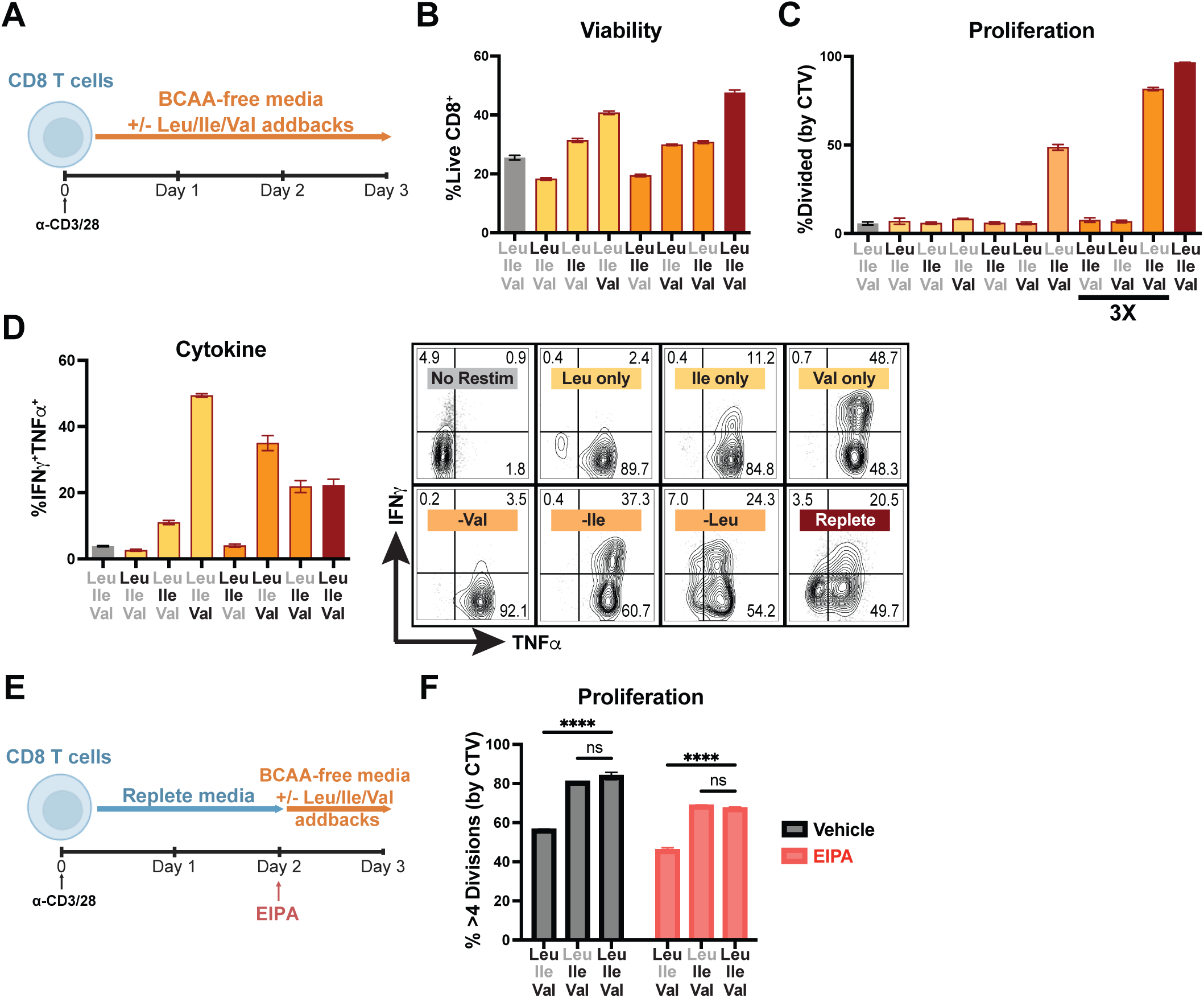
Isoleucine and Valine recover T cell division and cytokine production after multiple days in culture, even in the context of macropinocytosis inhibition. (A) Schematic of Day 0-3 BCAA restriction and addback culture. Bulk CD8 T cells were isolated and purified from spleens of WT mice, activated with anti-CD3 and anti-CD28 antibodies (2ug/mL each) for 3 days in BCAA-free TCM with addback of varying combinations of Leu, Ile and Val (0.8mM or 2.4mM [3X] each as indicated). (B) Frequency of viable CD8 T cells after Day 0-3 BCAA culture (2.4mM addbacks). (C) Frequency of dividing cells after Day 0-3 BCAA culture (% of cells with CTV dilution, CTV stain performed on Day 0). (D) Frequency and representative flow plots of cytokine-producing CD8 T cells after Day 0-3 BCAA culture (2.4mM addbacks). (E) Schematic of Day 2-3 BCAA restriction with EIPA and addback culture. Bulk CD8 T cells were isolated and purified from spleens of WT mice, activated with anti-CD3 and anti-CD28 antibodies (2ug/mL each) for 2 days in replete TCM followed by 1 day in BCAA-free TCM with addback of varying combinations of Leu, Ile and Val (0.8mM each) and simultaneous treatment with 25uM EIPA or vehicle control. (F) Frequency of highly dividing (>4 divisions) cells as measured by CTV dilution (CTV stain performed immediately after CD8 purification on Day 0 of culture). All error bars are representative of 3 technical replicates. Statistical significance in (F) was calculated using two-way ANOVA with multiple comparisons and Sidak’s correction. ****p<0.0001.

**Figure EV2:**
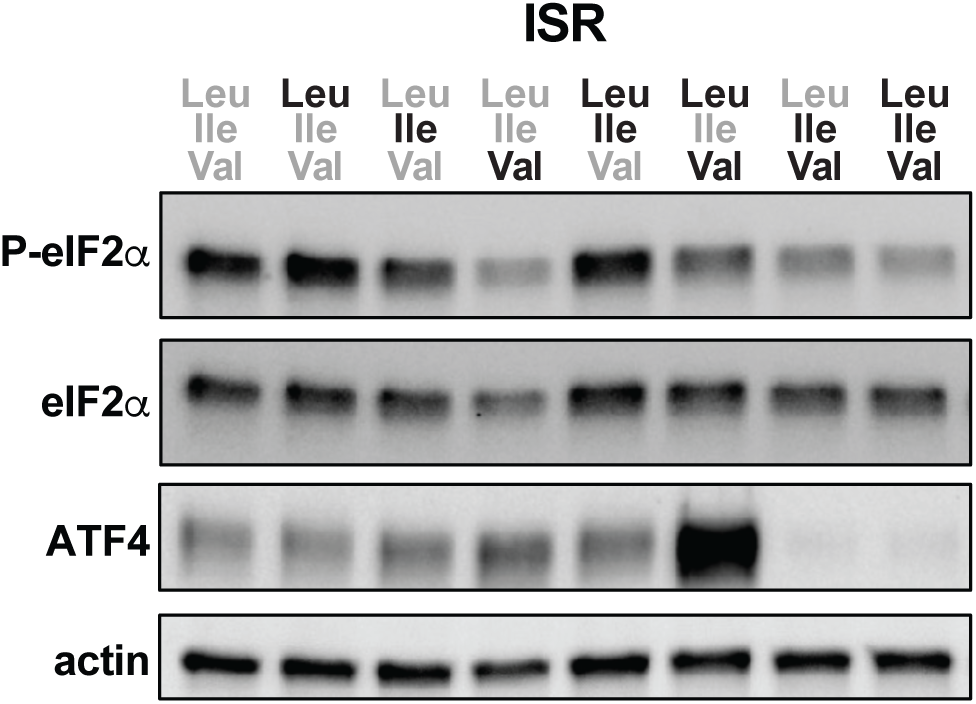
Differential sensing of individual BCAA availability by the integrated stress response. Western blot of integrated stress response signaling in acute BCAA culture as shown in 2A.

**Figure EV3:**
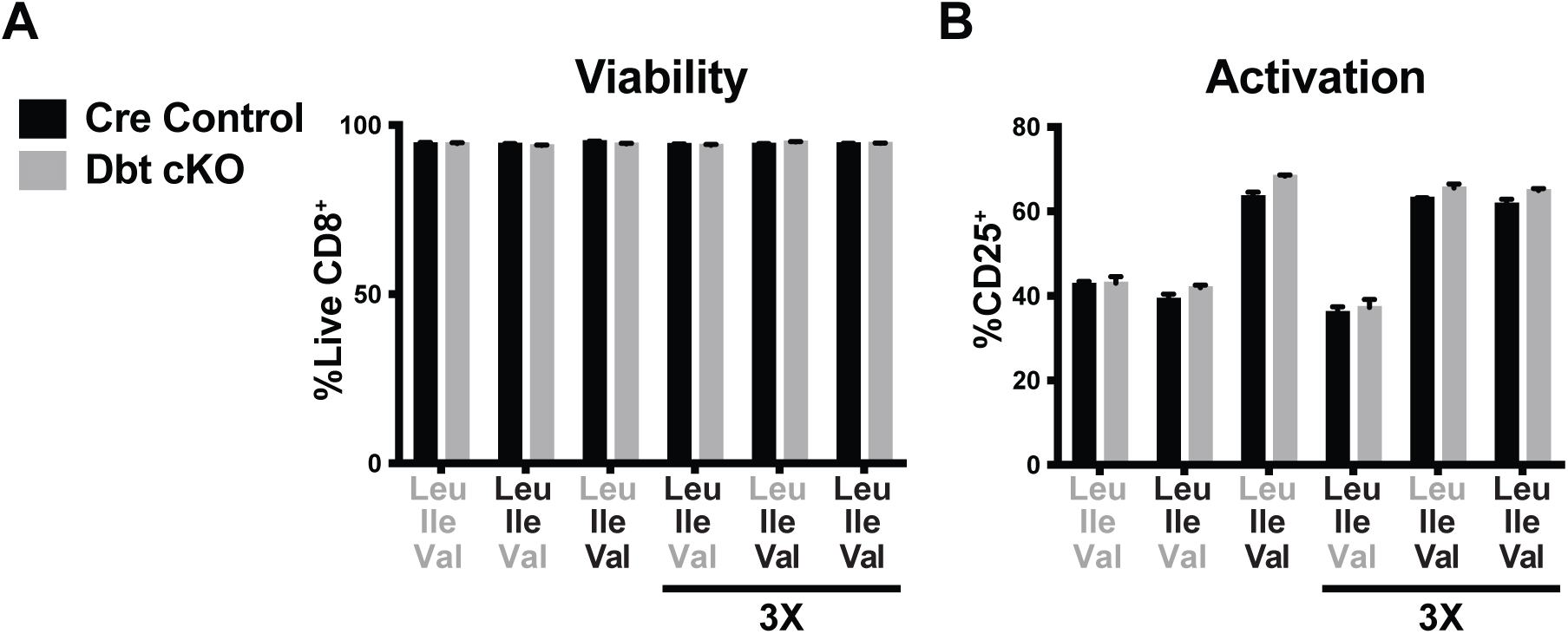
BCAA catabolism has negligible impact on CD8 T cell viability or Val-induced activation during one day of activation. (A,B) Frequency of viable CD8 T cells and activation (by CD25 expression induction) in Day 0-1 BCAA culture as described in (1A) (0.8mM or 2.4mM [3X] BCAA addbacks as indicated). All error bars are representative of 3 technical replicates.

**Figure EV4:**
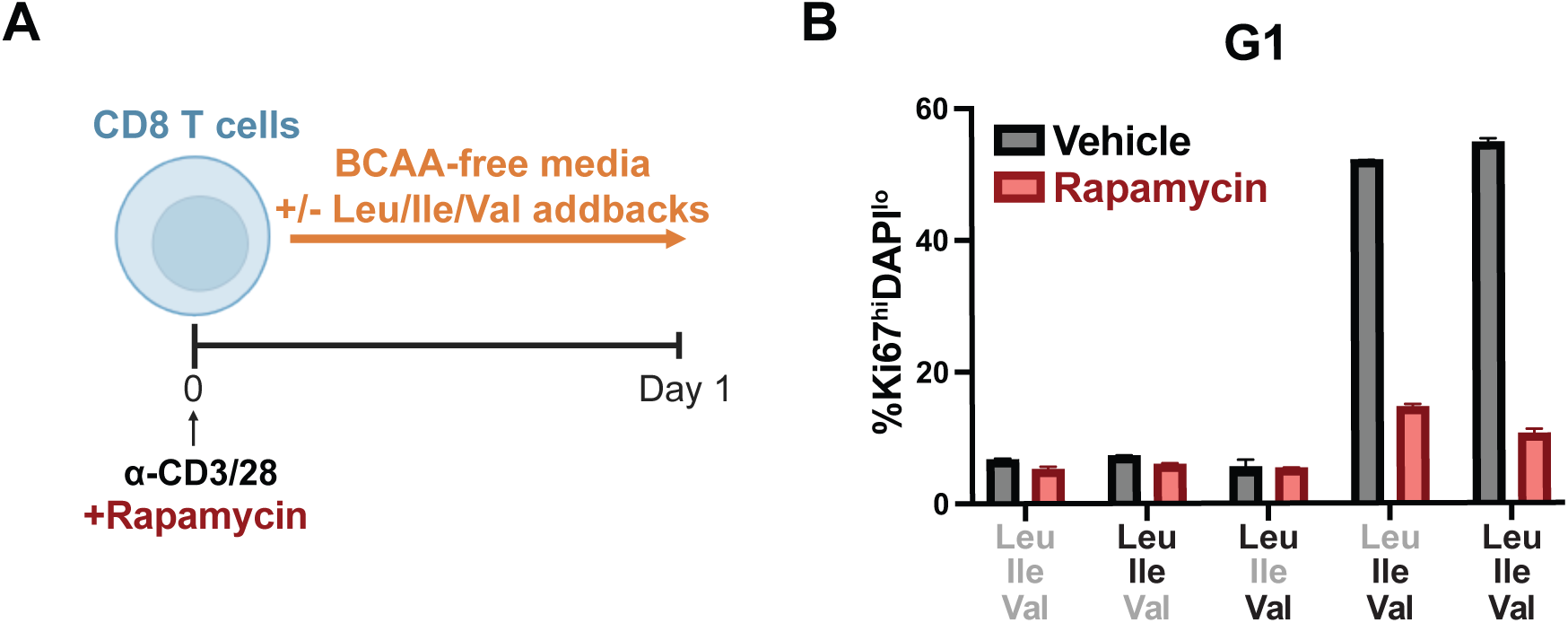
Ile and Val directed cell cycle entry during first day of activation is dependent on mTORC1. (A) Schematic of Day 0-1 BCAA restriction and addback culture with concurrent rapamycin treatment. Bulk CD8 T cells were isolated and purified from spleens of WT mice, activated with anti-CD3 and anti-CD28 antibodies (2ug/mL each) for 1 day in BCAA-free T cell media (TCM) with addback of varying combinations of Leu, Ile and Val (0.8mM each) and concurrent treatment with 50nM rapamycin or vehicle control. (B) Frequency of cells entering cell cycle after Day 0-1 BCAA culture with rapamycin. All error bars are representative of 3 technical replicates.

